# Dendrite-targeting inhibitory interneurons form biased circuits with deep and superficial pyramidal cells in hippocampal CA1

**DOI:** 10.1101/2025.06.04.657895

**Authors:** Aidan C. Johantges, Meretta A. Hanson, Alec H. Marshall, Alireza Safa, Emily K. Payne, Noor Bibi, Jason C. Wester

## Abstract

In CA1 hippocampus, pyramidal cells (PCs) can be classified as deep or superficial based on their radial position within the stratum pyramidale. Deep and superficial PCs form biased circuits with perisomatic-targeting PV+ basket cells, but it is unknown if such cell-type-specific circuit motifs extend to dendrite-targeting interneurons. Using male and female mice, we investigated synaptic connectivity and physiology in brain slices from four transgenic lines thought to capture distinct subsets of interneurons: SST-IRES-Cre, Nkx2.1-Cre, Chrna2-Cre, and Htr3a-GFP. First, we found that oriens-lacunosum moleculare (OLM) cells captured by the Chrna2-Cre line are a subset of Htr3a-GFP+ cells in the hippocampus. This novel finding is consistent with previous work showing Nkx2.1-Cre OLM cells are distinct from both Chrna2-Cre and Htr3a-GFP+ cells. Indeed, in paired whole-cell recordings, Nkx2.1-Cre+ interneurons in the stratum oriens, but not Chrna2-Cre+ or Htr3a-GFP+ cells, received more excitatory synaptic connections from superficial PCs relative to deep PCs. Next, we used optogenetic-assisted circuit mapping to investigate inhibition along the proximal and distal dendrites of PCs. We found that superficial PCs received stronger inhibition along their proximal dendrites than deep PCs from SST+ interneurons. Furthermore, this circuit motif was dependent on layer but not PC projection class. Finally, Chrna2-Cre OLM cells provided stronger inhibition to the distal dendrites of deep PCs relative to superficial PCs. Our data reveal that superficial and deep PCs engage in cell-type-specific circuits with dendrite-targeting interneurons. Furthermore, they support that Nkx2.1-Cre OLM cells and Chrna2-Cre/Htr3a-GFP OLM cells are distinct subtypes that form unique circuits in CA1.

## INTRODUCTION

The hippocampus is necessary for learning and memory and communicates with the rest of the brain via output from pyramidal cells in region CA1. These pyramidal cells receive excitatory synaptic input on their proximal dendrites from CA3 Schaffer collaterals and on their distal dendrites from the entorhinal cortex and thalamus (Chittajallu et al., 2017; Masurkar et al., 2017). This excitation is regulated by a diverse set of inhibitory interneurons that target distinct neuronal compartments (e.g., soma, proximal dendrites, and distal dendrites) (Klausberger and Somogyi, 2008; Pelkey et al., 2017). Furthermore, circuits with inhibitory interneurons play an important role in coordinating ensembles of pyramidal cells, as CA1 pyramidal cells rarely make excitatory synaptic connections between each other (Knowles and Schwartzkroin, 1981; Deuchars and Thomson, 1996).

Recent work found that CA1 pyramidal cells are heterogeneous and include multiple subtypes identified by molecular expression profile, electrophysiological properties, and morphology (Cembrowski et al., 2016; D’Amour et al., 2020; Masurkar et al., 2020; Hanson et al., 2025). These differences correlate with pyramidal cell position along the radial axis, leading to the hypothesis that the stratum pyramidale contains distinct superficial and deep sublayers that encode unique information and provide separate output channels from CA1 (Slomianka et al., 2011; Soltesz and Losonczy, 2018). Indeed, superficial and deep pyramidal cells encode unique spatial information during navigation tasks and are differentially recruited during sharp wave ripples (Valero et al., 2015; Danielson et al., 2016; Geiller et al., 2017; Sharif et al., 2021; Esparza et al., 2025). Cell-type-specific circuits with inhibitory interneurons importantly contribute to these laminar functional differences. Work from multiple labs found that deep pyramidal cells receive more inhibitory synapses from parvalbumin (PV)-expressing basket cells than superficial pyramidal cells, which results in biased feedforward inhibition of CA3 Schaffer collateral input (Lee et al., 2014; Valero et al., 2015; Masurkar et al., 2017; Hanson et al., 2025). Furthermore, superficial pyramidal cells provide greater excitatory drive to PV basket cells, which may allow them to indirectly regulate excitation of deep pyramidal cells (Lee et al., 2014; Hanson et al., 2025).

It is unknown whether layer-dependent circuit motifs exist for other interneuron subtypes. These include somatostatin (SST)-expressing interneurons whose cell bodies principally reside in or near the stratum oriens (S.O.) (Müller and Remy, 2014; Pelkey et al., 2017; Agmon and Barth, 2024; Chamberland et al., 2024). SST+ interneurons are diverse and include bistratified cells (BiCs) that target proximal and basal dendrites and oriens-lacunosum moleculare (OLM) cells that target distal dendrites. Furthermore, there is evidence that OLM cells include distinct subtypes, which are differentially captured by transgenic mouse lines expressing Cre or GFP under the control of promotors for *Nkx2.1*, *Htr3a*, and *Chrna2* (Chittajallu et al., 2013; Winterer et al., 2019; Chamberland et al., 2024). However, OLM cells from all three lines have similar morphologies and electrophysiological properties. Furthermore, transcriptomic differences between them are subtle (Winterer et al., 2019). Thus, the extent to which these OLM cells engage in functionally distinct circuits remains unclear. A recent study found differential gene expression between Nkx2.1-Cre+ and Chrna2-Cre+ OLM cells (Chamberland et al., 2024). Furthermore, it found that OLM cells represented in the Chrna2-Cre line target neighboring PV+ basket cells, while OLM cells from the Nkx2.1-Cre line largely do not (Chamberland et al., 2024). However, it is unknown if OLM cells captured by these different transgenic lines engage in cell-type-specific circuits with pyramidal cells.

Here, we used four different transgenic mouse lines, SST-IRES-Cre, Nkx2.1-Cre, Chrna2-Cre, and Htr3a-GFP, to investigate circuits between dendrite-targeting inhibitory interneurons and deep and superficial pyramidal cells in hippocampal CA1. First, we show that OLM cells captured by the Chrna2-Cre mouse line are a subset of Htr3a-GFP+ interneurons in the hippocampus. Next, we show that superficial and deep pyramidal cells form cell-type-specific circuits with both OLM cells and interneurons that target proximal dendrites, such as BiCs. Furthermore, OLM circuit motifs are unique for those captured by the Nkx2.1-Cre or Chrna2-Cre mouse lines. Thus, our data provide novel insight into cell-type-specific circuits that regulate dendritic inhibition in CA1.

## RESULTS

We investigated local circuits between pyramidal cells and dendrite-targeting SST+ inhibitory interneurons in CA1 hippocampus. To target different subsets of SST+ interneurons we used four transgenic mouse lines: SST-IRES-Cre, Nkx2.1-Cre, Htr3a-GFP, and Chrna2-Cre. The three Cre driver lines were further crossed to the floxed tdTomato reporter line, Ai14 (Madisen et al., 2010) (**Fig. 1A**). The SST-IRES-Cre line (Taniguchi et al., 2011) broadly captures SST+ interneurons, which are concentrated in the stratum oriens (**Fig. 1Ai**). However, a small number of excitatory pyramidal cells also expressed Cre, consistent with previous work (Mikulovic et al., 2015) (**Fig. 1Ai**; two pyramidal cells are visible on far left). The Nkx2.1-Cre line captures inhibitory interneurons derived from the medial ganglionic eminence (MGE) during embryogenesis (Xu et al., 2008), which includes SST+ and PV+ subtypes (Miyoshi et al., 2007; Tricoire et al., 2011) (**Fig. 1Aii**). The Htr3a-GFP line is a BAC transgenic that captures interneurons derived from the caudal ganglionic eminence (CGE) that are concentrated in the strata pyramidale, radiatum, and lacunosum moleculare and primarily express cholecystokinin (CCK), calretinin (CR), vasoactive intestinal peptide (VIP), and reelin (but are SST-negative) (Lee et al., 2010; Tricoire et al., 2011) (**Fig. 1Aiii**). However, this line also captures SST+ cells in the S.O. (including OLM cells) that are non-overlapping with those captured by the Nkx2.1-Cre line (Chittajallu et al., 2013). Finally, the Chrna2-Cre line captures a specific subset of OLM cells that express the nicotinic acetylcholine receptor α2, and are referred to as OLMα2 cells (Leao et al., 2012; Siwani et al., 2018) (**Fig. 1Aiv**). To map circuits in CA1, we targeted interneurons from these four transgenic lines in the stratum oriens for paired whole-cell recordings with deep and superficial pyramidal cells. We also used the SST-IRES-Cre and Chrna2-Cre lines for optogenetic-assisted circuit mapping.

**Figure 1.**
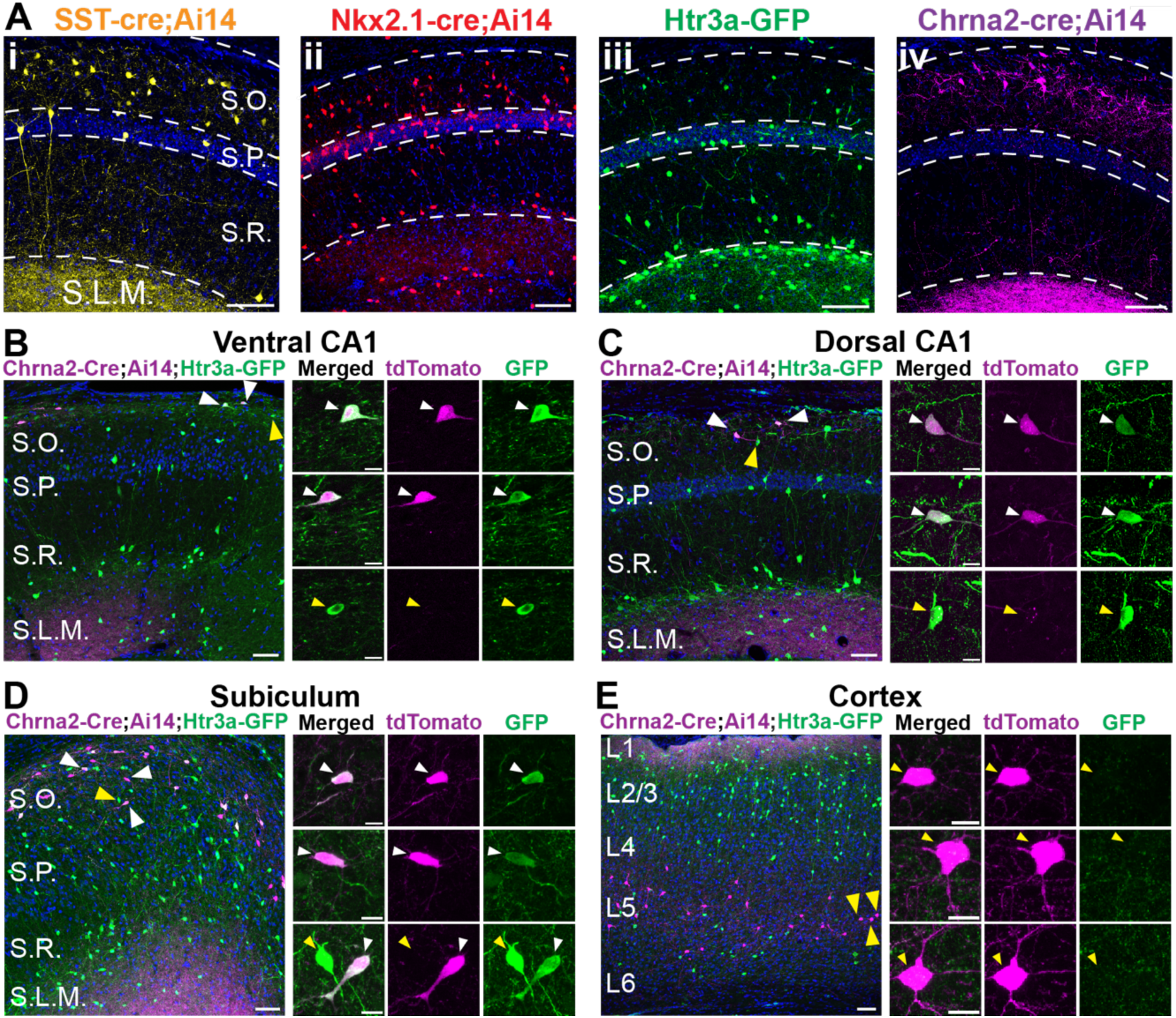
Transgenic mouse lines to capture subpopulations of SST+ interneurons in the stratum oriens. **A)** Examples of fluorescent protein expression in dorsal CA1 in transgenic mice used in this study. (i) SST-IRES-Cre;Ai14 labels SST+ cells primarily within the S.O. and S.R., but also some pyramidal cells (pictured on the left). (ii) Nkx2.1-Cre;Ai14 labels SST+ and PV+ interneurons throughout the hippocampus. (iii) Htr3a-GFP primarily labels VIP+, CR+, and CCK+ interneurons in the S.P., S.R., and S.L.M, but also a subset of SST+ interneurons within the S.O. (iv) Chrna2-Cre;Ai14 labels a subpopulation of OLM cells within the S.O. S.O, stratum oriens; S.P., stratum pyramidale; S.R., stratum radiatum; S.L.M., stratum lacunosum-moleculare; scale bars = 100 µm. **B)** In horizontal sections of ventral CA1, Chrna2-Cre+ cells (identified by tdTomato expression via Cre recombination) are common. Three example cells are shown at high magnification on the right. All Chrna2-Cre+ cells are Htr3a-GFP+ (white arrows), but not all Htr3a-GFP+ interneurons are Chrna2-Cre+ (yellow arrows). **C)** In coronal sections of dorsal CA1, Chrna2-Cre+ cells are sparse, but all are Htr3a-GFP+ (white arrows). **D)** In horizontal sections of ventral subiculum, Chrna2-Cre+ cells are common. Like in CA1, all Chrna2-Cre+ cells are Htr3a-GFP+ (white arrows), but not all Htr3a-GFP+ interneurons are Chrna2-Cre+ (yellow arrows). **E)** In neocortex, Chrna2-Cre+ cells are concentrated in layer 5. In contrast to CA1 and subiculum, no Chrna2+ cells co-express Htr3a-GFP (yellow arrows). For (**B – E**), scale bars for low magnification images on the left are 50 µm, and scale bars for high magnification images on the right are 10 µm.

### OLMα2 cells are Htr3a-GFP+ interneurons in the hippocampus and subiculum but not the neocortex

The Nkx2.1-Cre and Htr3a-GFP mouse lines label non-overlapping sets of inhibitory interneurons in the S.O. of CA1 (Chittajallu et al., 2013). Both lines capture OLM cells, but there is debate regarding whether these are separate subtypes (Asgarian et al., 2019; Winterer et al., 2019). Furthermore, it is unknown if the OLMα2 cells captured by the Chrna2-Cre line are a subset OLM cells captured by either of these transgenics. Thus, the extent to which OLM cells captured by these different transgenic lines should be considered as distinct types is unclear. To investigate overlap between these mouse lines, we generated Chrna2-Cre;Ai14;Htr3a-GFP mice. Strikingly, we found that all Chrna2-Cre-tdTomato+ cells were also GFP+ in both ventral and dorsal CA1 hippocampus (**Figs. 1B and 1C**) and in the subiculum (**Fig. 1D**). Thus, in these structures, OLMα2 cells are a subset of Htr3a-GFP+ cells. However, we observed no overlap in the neocortex (**Fig. 1E**). Although this was surprising, it is consistent with previous findings that SST-expression in Htr3a-GFP+ interneurons is very rare in the neocortex relative to hippocampus (Lee et al., 2010; Vucurovic et al., 2010; Chittajallu et al., 2013). We conclude that the previously reported subpopulation of Htr3a-GFP+ OLM cells in CA1 are OLMα2 cells (Leao et al., 2012; Chittajallu et al., 2013). Furthermore, these data provide additional evidence that OLMα2/Htr3a-GFP+ OLM cells are distinct from those captured in the Nkx2.1-Cre line (Chittajallu et al., 2013; Chamberland et al., 2024).

### S.O. interneurons from the Nkx2.1-lineage receive biased excitatory input from superficial pyramidal cells

To investigate circuits, we began by assaying unitary excitatory synaptic connections from pyramidal cells to interneurons in the S.O., which can be observed using paired-whole cell recordings. We targeted a deep or superficial pyramidal cell, and a fluorescent interneuron tagged with either tdTomato via Cre recombination or GFP (**Fig. 2A**). We performed 247 paired recordings, including approximately 60 in each transgenic mouse line, evenly split between deep and superficial pyramidal cells (**Fig. 2B**). We first analyzed the probability of finding excitatory synaptic connections from deep or superficial pyramidal cells to S.O. interneurons. In data pooled from all connections tested, we found a strong bias in the connectivity rate from superficial pyramidal cells (superficial: 25% (31/125) vs. deep: 10% (12/122); **Fig. 2B, left**). Next, we considered the contribution of each individual transgenic line to these connection probabilities. The connectivity rate bias was evident for interneurons labeled in the SST-IRES-Cre;Ai14 line and was strongest among those labeled in the Nkx2.1-Cre;Ai14 line (**Fig. 2B, right**). However, surprisingly, the bias was absent when interneurons were targeted in either the Htr3a-GFP or Chrna2-Cre;Ai14 lines (**Fig. 2B, right**). Thus, superficial and deep pyramidal cells differentially target inhibitory interneurons in the S.O. However, this circuit motif also depends on the subtypes of the interneurons, which are differentially represented among Nkx2.1-Cre+ and Chrna2-Cre+/Htr3a-GFP+ cells.

**Figure 2.**
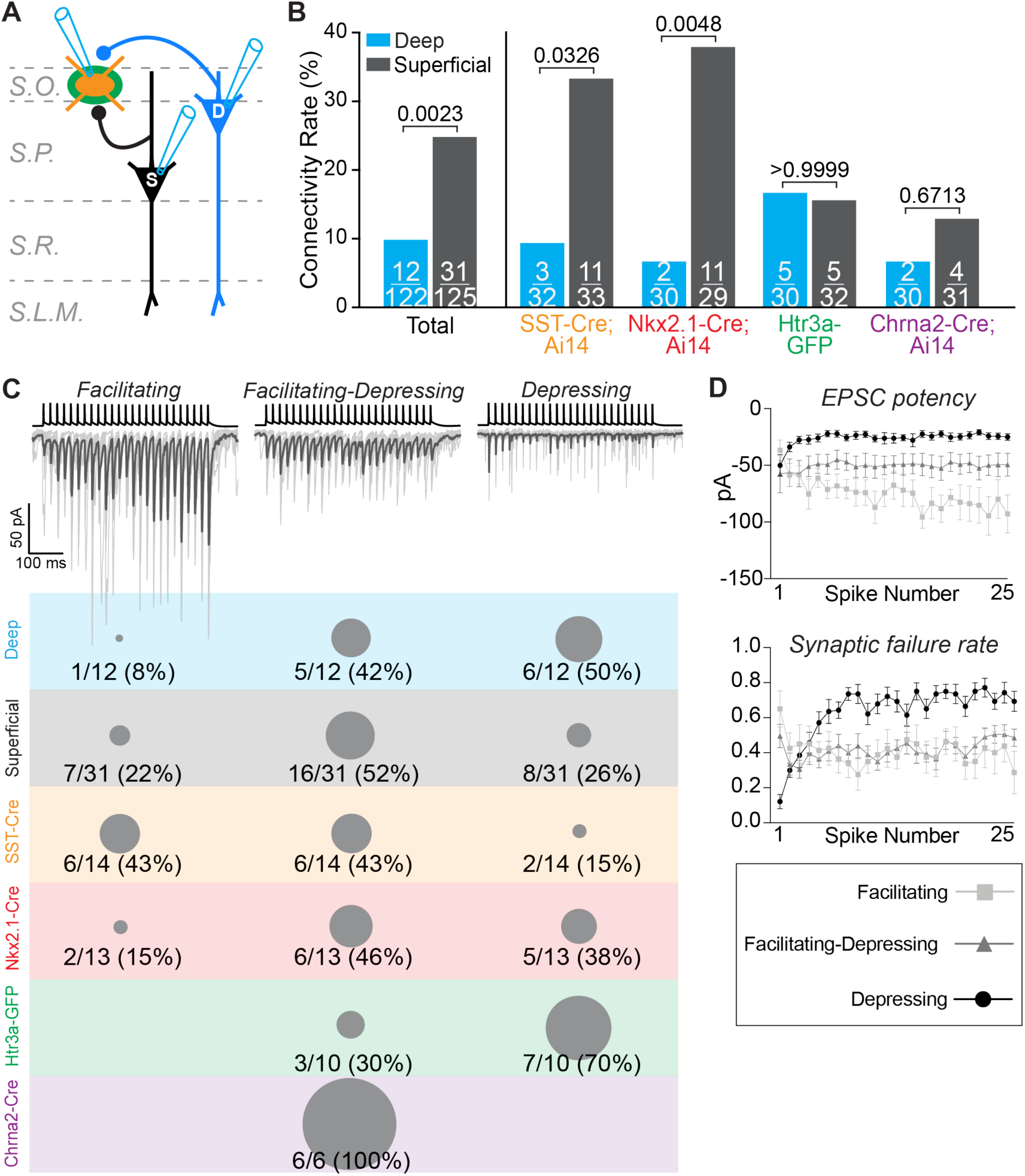
The connectivity rate and physiology of excitatory synapses from pyramidal cells to inhibitory interneurons in the stratum oriens are cell-type-dependent. **A)** Experimental configuration for paired whole-cell patch clamp recordings to investigate the connectivity and physiology of excitatory synapses from deep and superficial pyramidal cells to interneurons in the S.O. **B)** Total numbers of synaptic connections observed per number of tests performed for deep and superficial pyramidal cells in each transgenic mouse line. Data on the left are pooled from all connections tested and found. Superficial pyramidal cells provide biased excitatory synaptic connections to interneurons in the S.O. This bias is observed for SST-IRES-Cre+ and Nkx2-1-Cre+ interneurons but not Htr3a-GFP+ or Chrna2-Cre+ interneurons. SST-IRES-Cre;Ai14: n = 65 paired recordings in 5 mice; Nkx2.1-Cre;Ai14: n = 59 paired recordings in 14 mice; Htr3a-GFP: n = 62 paired recordings in 16 mice; Chrna2-Cre n = 61 paired recordings in 5 mice. The numbers of Nkx2.1-Cre;A14 and Htr3a-GFP mice are higher because additional interneurons were recorded individually in slices from these mice to support cell-type identification via unbiased clustering in Figure 4. P values from Fisher’s exact test. **C)** (*top*) Representative examples of the three types of short-term plasticity observed at synapses from pyramidal cells (presynaptic action potentials in black) to interneurons in the S.O. (postsynaptic currents in gray). Synaptic responses from individual trials are in light gray; dark gray traces are the averages of 10 single trials. (*bottom*) Proportions of each postsynaptic response type separated by 1) connections from all deep pyramidal cells (blue), 2) from all superficial pyramidal cells (gray), and 3) all synaptic connections observed in each transgenic mouse line (bottom colors). The gray dots represent proportions relative to 100% (see Chrna2-Cre, bottom purple). **D)** Quantification of excitatory postsynaptic current (EPSC) potency (the average postsynaptic current evoked by each presynaptic spike over 10 trials, excluding synaptic failures) and synaptic failure rate (percentage of trials in which a presynaptic action potential did not evoke a postsynaptic current). This analysis shows that synapses demonstrating facilitation, facilitation-depression, and depression are quantitatively distinct. N = 8 facilitating synapses, n = 20 facilitating-depressing synapses, and n = 14 depressing synapses.

Next, we analyzed the physiology of the postsynaptic excitatory connections observed in our paired recordings. We evoked trains of presynaptic action potentials in pyramidal cells, which allowed us to investigate the short-term postsynaptic dynamics of excitatory inputs to S.O. interneurons (**Fig. 2C**). We observed three types of excitatory postsynaptic responses, which were consistent with previous studies of OLM and bistratified cells in CA1 S.O.: facilitating, facilitating-depressing, and depressing (Ali et al., 1998; Ali and Thomson, 1998; Losonczy et al., 2002) (**Fig. 2C, top**). These responses were qualitatively distinct (**Fig. 2C, top**) and could be identified quantitatively based on synaptic potency (average synaptic current amplitude for trials in which a presynaptic action potential evoked vesicle release) and synaptic failure rate (percentage of trials in which a presynaptic action potential did not evoke vesicle release) (Stevens and Wang, 1995) (**Fig. 2D**). We found that facilitating and facilitating-depressing synapses were most common from superficial pyramidal cells and were the majority observed in Nkx2.1-Cre;Ai14 mice (**Fig. 2C, bottom, gray and red**). Conversely, depressing synapses were most common from deep cells and were the majority observed in Htr3a-GFP mice (**Fig. 2C, bottom, blue and green**). Interestingly, all excitatory synaptic connections from pyramidal cells to OLMα2 cells were facilitating-depressing (**Fig. 2C, bottom, purple**). Thus, excitatory synaptic dynamics depend on both pyramidal cell and S.O. interneuron subtype identified in each transgenic.

Interneurons in CA1 S.O. are diverse and include subtypes that differentially target pyramidal cell dendrites in the strata oriens, radiatum, and lacunosum moleculare (Tricoire et al., 2011). Thus, we next asked if the bias in connectivity rate from superficial pyramidal cells (**Fig. 2B**) could be attributed to a specific subtype of these interneurons. To parse interneurons into different types, we collected detailed data on intrinsic membrane properties from 189 cells and filled them with biocytin during recording for post-hoc morphological analysis (**Fig. 3**). These included interneurons sampled during paired whole-cell recordings and individually from all four transgenic lines (31 cells from SST-IRES-Cre, 73 from Nkx2.1-Cre, 25 from Chrna2-Cre, and 60 from Htr3a-GFP). We measured passive membrane properties (**Figs. 3A – C**), details of individual action potentials (**Fig. 3D**), and firing rate dynamics during steps of current injection (**Figs. 3E – G**). We also recorded the resting membrane potential prior to injecting bias current to hold cells at -70 mV (not shown). Finally, we classified cells based on recovered morphology, focusing on axonal arborization across CA1 strata. We broadly parsed cells into three categories: OLM cell, non-OLM cell, and incomplete morphology (**Fig. 3H**). To classify cells as OLM, we required recoveries to include preserved axon in the stratum lacunosum moleculare; this allowed us to distinguish them from recently described hippocampal-septal cells, which strongly resemble OLM cells but whose axons stop within the stratum radiatum (Takács et al., 2024). Thus, non-OLM cells include bistratified cells (BiCs), hippocampal-septal cells, and undefined subtypes.

**Figure 3.**
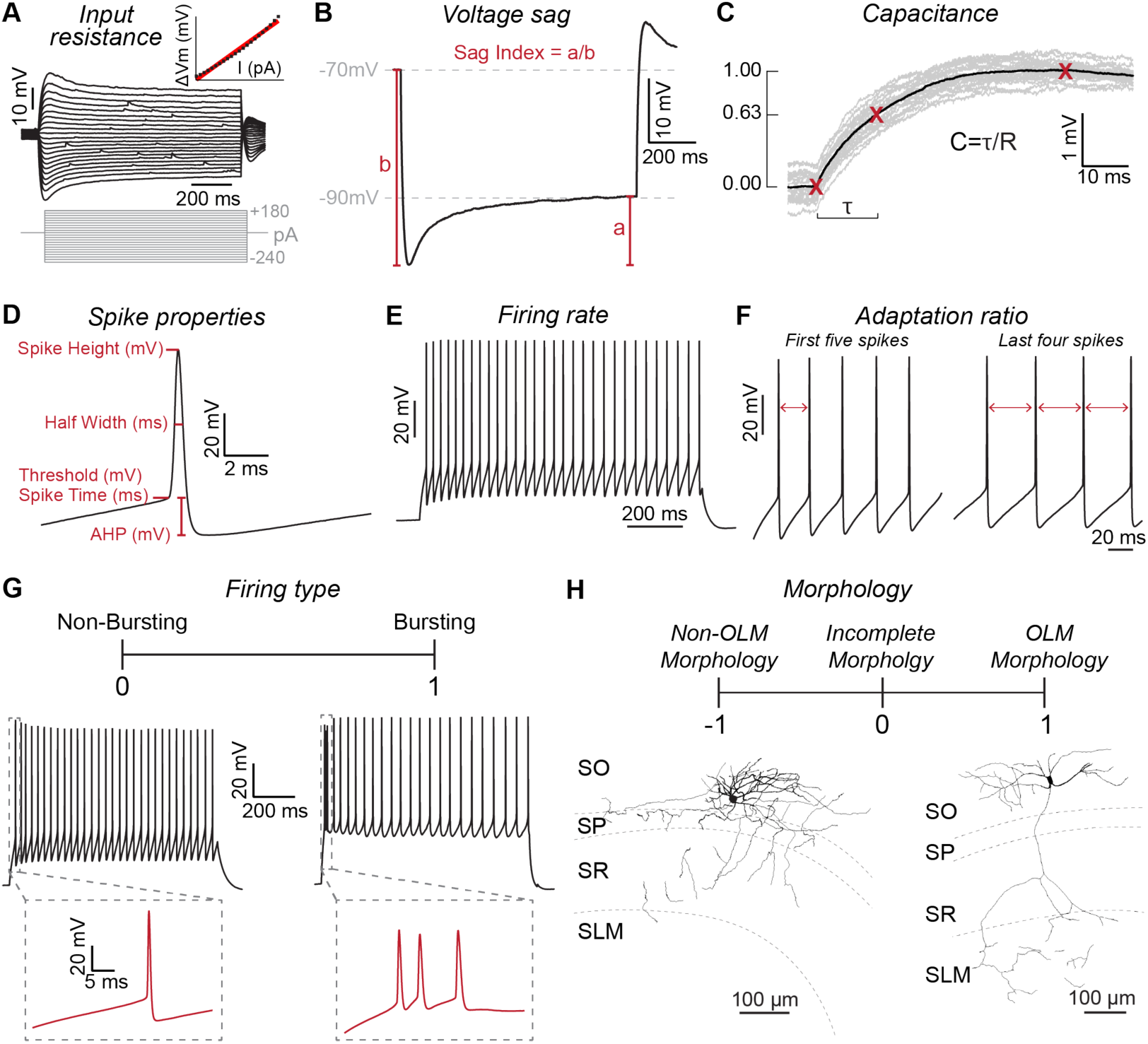
Physiological and morphological properties used for K-means clustering of interneurons. **A)** Input resistance was measured by linear regression of voltage deflections in response to constant amplitude current steps. **B)** Some stratum oriens interneurons exhibit robust voltage sag in response to hyperpolarizing current injections. To quantify voltage sag, we injected a current step of sufficient amplitude for the membrane potential to settle at -90 mV (value a). To calculate the sag index, we took the ratio of this value (a) relative to the peak membrane potential hyperpolarization measured at the start of the injected current step (value b). **C)** The time constant (τ) and membrane capacitance of each interneuron were calculated by injecting 10 pA current pulses to minimize activation of voltage-dependent conductances. **D)** The minimum current amplitude was injected to evoke at least two action potentials during a step pulse. For both action potentials, we measured voltage threshold and time to threshold, spike height, spike half width at half height, and hyperpolarization voltage relative to voltage threshold. **E)** Firing rate was measured during current pulses at twice the rheobase amplitude. **F)** To measure firing rate adaption during current pulses at twice the rheobase amplitude, we took the ratio of the time between the first two action potentials to the average time between the final four action potentials. **G)** During current step injections, we identified interneurons that fire initial action potential bursts (*right*) or singlets (*left*). For principal component analysis and K-means clustering, bursting cells were assigned a value of 1 and non-bursting cells were assigned a value of 0. **H)** We classified interneurons by their morphology recovered post-hoc after filling with biocytin during recording. Cells with horizontally-projecting dendrites and a primary axon that arborized within the S.L.M. were classified as OLM cells and assigned the value of 1. Cells with complete morphologies that lacked a primary axon arborizing within S.L.M were identified as non-OLM and assigned a value of -1. Cells without a complete morphology were assigned a value of 0.

We used these data (**Fig. 3**) to perform principal component analysis and unsupervised K-means clustering to segregate S.O. interneurons into groups (**Fig. 4**). From principal component analysis we found that properties of individual action potentials, voltage sag, and input resistance primarily contributed to distinguishing cells (**Fig. 4A**). Using NbClust (Charrad et al., 2014), we determined that the optimal number of clusters for K-means analysis was 3 (**Fig. 4B**). We plotted the results from principal component and K-means analysis, which revealed three well-segregated clusters of interneurons (**Fig. 4C**). Next, we analyzed the proportion of interneurons from each cluster recorded in the four different transgenic mouse lines (**Fig. 4D**). Most interneurons recorded in the Chrna2-Cre;Ai14 line were from Cluster 1 (**Fig. 4D**). Importantly, all OLM cells share similar electrophysiological and morphological properties (Chittajallu et al., 2013; Winterer et al., 2019; Chamberland et al., 2024), thus, Cluster 1 likely represents OLM cells recorded in the other three transgenic lines. Interneurons recorded in the Nkx2.1-Cre;Ai14 line were primarily from Clusters 1 and 3, while those in the Htr3a-GFP mouse line were evenly distributed among all three clusters. Finally, most interneurons in the SST-IRES-Cre;Ai14 line were from Cluster 1 (**Fig. 4D**). Next, we matched the interneuron cluster number to the type of short-term excitatory postsynaptic dynamics observed in paired recordings (**Fig. 4E**). Interneurons in Cluster 1 received mostly facilitating or facilitating-depressing synapses, those in Cluster 2 received only facilitating-depressing synapses, and those in Cluster 3 received only depressing synapses (**Fig. 4E**). These data are consistent with previous work showing that excitatory synapses to OLM cells are either facilitating or facilitating-depressing, while those to bistratified cells include synaptic depression (Ali et al., 1998; Ali and Thomson, 1998; Losonczy et al., 2002). It is also consistent with our finding that all excitatory synapses to OLMα2 cells are facilitating-depressing (**Fig. 2C**). Thus, Clusters 1 and 2 represent primarily OLM cells, while Cluster 3 primarily represents bistratified cells and other cell-types that target proximal dendrites. Finally, our clusters are consistent with previous work that used principal component analysis and K-means clustering to separate OLM and bistratified cells into separate clusters based on their electrophysiological properties (Chamberland et al., 2024).

**Figure 4.**
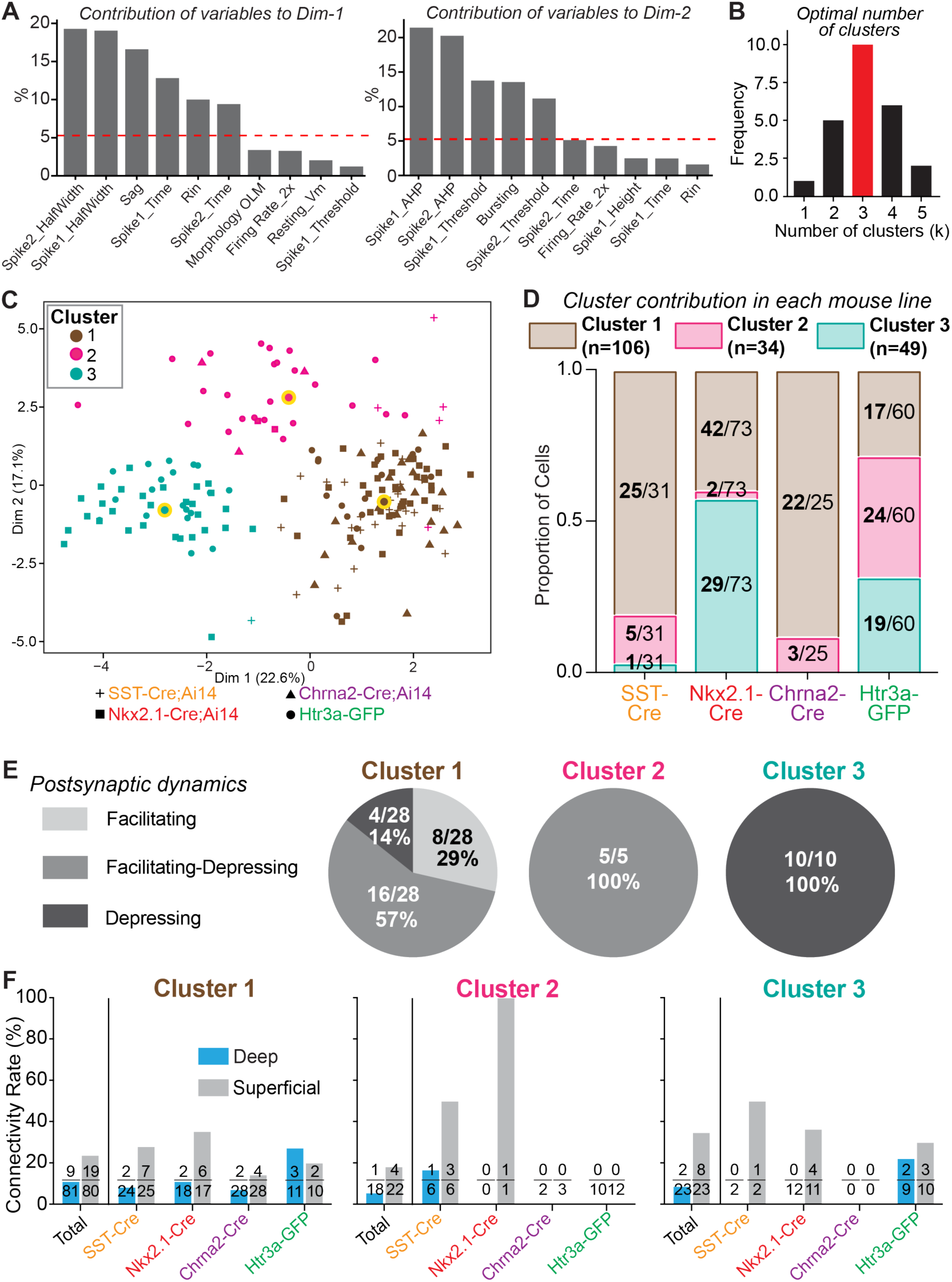
Superficial pyramidal cells provide biased synaptic connections to all subtypes of Nkx2.1-Cre+ interneurons in the S.O. **A)** Top 10 variables from principal component analysis (PCA) that distinguish interneuron subtypes. The dotted red lines denote the expected average contribution of each variable if it were uniform. The first six variables of each dimension explain over 80% of variability of that dimension. **B)** The optimal number of clusters indicated by NbClust was k = 3 and thus used for further analysis. **C)** Principal component analysis and K-means clustering plot of interneuron properties (*see* Figure 3). Each cell has a unique icon that corresponds to the genotype of the mouse from which it was recorded (*bottom*). Cluster 1, n = 106 cells; Cluster 2, n = 34 cells; Cluster 3, n = 49 cells. Cluster centroids are circled in yellow. **D)** For each transgenic mouse line from which interneurons were recorded, we plotted the proportion of interneurons from each cluster. Note that nearly all interneurons recorded in slices from the Chrna2-Cre mouse line are in Cluster 1. Thus, we infer that Cluster 1 represents OLM cells in all mouse lines. Cluster 3 likely contains subtypes that include bistratified cells. **E)** We identified the cluster assignment of each interneuron for which we observed a unitary synaptic connection from a pyramidal cell. Interneurons in Cluster 1 primarily received facilitating or facilitating-depressing synapses, and interneurons in Cluster 2 only received facilitating-depressing synapses. This is consistent with our finding that Chrna2-Cre+ cells are only found in Cluster 1 and Cluster 2. In contrast, all interneurons in Cluster 3 receive depressing synapses, which is consistent with our conclusion that these are likely bistratified cells and related subtypes. **F)** Synaptic connectivity rate data from paired whole-cell recordings between pyramidal cells and interneurons were reanalyzed according to the cluster assignment of each postsynaptic interneuron tested. This analysis revealed that superficial pyramidal cells provide biased excitatory input to all subtypes off Nkx2.1-Cre+ interneurons. However, there is no bias in connectivity rate from pyramidal cells to Htra3-GFP+ interneurons of any subtype.

Based on this clustering, we reanalyzed our data on synaptic connectivity rates and parsed cells according to both cluster number and transgenic mouse line. Strikingly, we found that superficial pyramidal cells provided biased synaptic connections to interneurons in all three clusters (**Fig. 4F**, left, total connections tested). Furthermore, in all three clusters this bias was strong for Nkx2.1-Cre+ and SST-IRES-Cre+ interneurons but not Chrna2-Cre+ or Htr3a-GFP+ interneurons (**Fig. 4F**). The connectivity rate data are similar for recordings made in the Nkx2.1-Cre and SST-IRES-Cre mouse lines because 70% of SST+ interneurons are captured by the Nkx2.1-Cre line in CA1 (Chittajallu et al., 2013). In summary, these data show that a subset of SST+ interneurons in the S.O. receive preferential excitatory drive from superficial pyramidal cells. These interneurons are selectively captured by the Nkx2.1-Cre line and include diverse subtypes that span OLM and bistratified cells. In contrast, OLMα2 cells captured by the Chrna2-Cre and Htr3a-GFP lines, and bistratified cells captured by the Htr3a-GFP line, receive fewer synaptic inputs from pyramidal cells, and their connectivity rates are comparable between superficial and deep cells.

### Superficial pyramidal cells receive greater inhibition along their proximal dendrites compared to deep pyramidal cells

Next, we investigated inhibitory synaptic connections from SST+ interneurons to superficial and deep pyramidal cells using channelrhodpsin-2 (ChR2)-assisted circuit mapping. To express ChR2 broadly in SST+ interneurons, we crossed SST-IRES-Cre mice to the Ai32 line, which encodes a Cre-dependent ChR2-EYFP fusion protein (Madisen et al., 2012). For these experiments we isolated inhibitory synapses by bath applying AMPA and NMDA receptor blockers and voltage-clamping pyramidal cells at 0 mV to render GABAergic currents outward. To focally activate ChR2, we delivered light pulses (2 ms duration) in trains of 10 at 10 Hz through a 40X water immersion objective.

We first focused the light on the stratum radiatum to activate inputs to the proximal region of pyramidal cell apical dendrites (**Fig. 5A**). We recorded superficial and deep pyramidal cells in pairs to allow us to perform direct comparisons of inhibitory input strength (**Figs. 5A and 5B**). Strikingly, ChR2-evoked responses in the stratum radiatum were consistently larger in amplitude in superficial cells relative to deep cells (**Fig. 5C**). Indeed, across the population, evoked inhibitory inputs to superficial cells were larger in most paired recordings (**Fig. 5D**), and average inhibitory currents were larger throughout the light pulse train (**Fig. 5E**). Because these synapses exhibited synaptic depression, for each pair we normalized the amplitude of the current evoked in the deep cell to that evoked in the simultaneously recorded superficial for each light pulse. This revealed that the inhibitory currents recorded in each pair scaled consistently throughout the train (**Fig. 5F**). Finally, we found no difference in the paired pulse ratio of inhibitory inputs to superficial and deep cells, suggesting comparable presynaptic vesicle release probability (**Fig. 5G**). Combined with our data on excitatory connections from paired whole-cell recordings (**Figs. 2 – 4**), these data suggest that superficial pyramidal cells engage in a feedback inhibitory circuit with SST+ interneurons that target proximal dendrites, which include BiCs (**Fig. 5H**).

**Figure 5.**
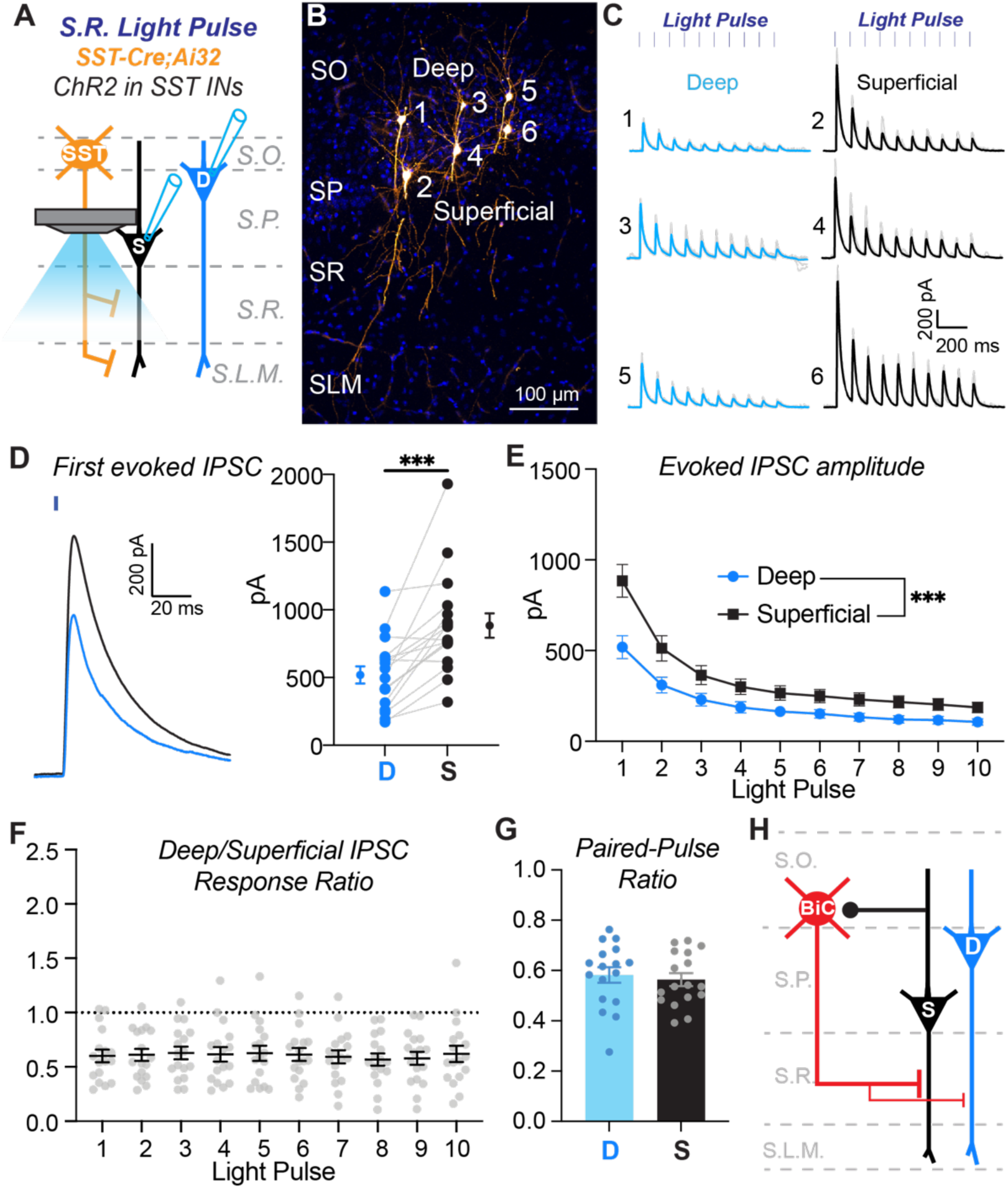
SST+ interneurons provide stronger inhibition to the proximal dendrites of superficial pyramidal cells relative to deep pyramidal cells. **A)** Experimental configuration to assay the strength of inhibitory input from SST+ interneurons to the proximal dendrites of superficial and deep pyramidal cells. Light was focused on the stratum radiatum (S.R.) to excite SST+ interneuron axons. Pairs of superficial and deep pyramidal cells were recorded simultaneously during light stimulation to directly compare inhibitory inputs. **B)** Representative example of a slice with recovered pyramidal cell bodies to demonstrate paired recordings of deep and superficial pyramidal cells in CA1. Three pairs were recorded in this slice (numbers 1+2, 3+4, and 5+6). Note, the tissue was re-sectioned for cell recovery and slide mounting, thus the full dendritic tree is not shown in this section. **C)** ChR2-evoked inhibitory currents recorded from the deep and superficial pyramidal cell pairs shown in (B). Blue and black lines are the average responses from 10 trials; light grey lines are responses from individual trials. The numbers relate to the corresponding cell body locations in (B). **D)** Average amplitudes of responses to the first light pulse in the train for the population of recordings. Gray lines connect cells recorded as a pair. At left is an expanded example of ChR2-evoked responses to the first light pulse in a deep (blue) and superficial (black) pyramidal cell pair. n = 17 paired recordings from n = 3 mice. *** p = 0.0001, paired t test (t(16) = 4.950). **E)** Average ChR2-evoked current amplitudes for the population of responses recorded in deep and superficial PCs during the light pulse train. N’s equivalent to (D). Main effect of cell type, *** p = 0.0003, repeated-measures two-way ANOVA: (F(1,16)= 20.65). **F)** Ratios of the average peak current amplitudes between deep and superficial PCs for the population. Ratio of deep/superficial PC response to 10 optogenetic light pulses. N’s equivalent to (D). Dotted grey line is the assumed value if the ratio of the responses were equivalent. Response ratios are lower than one, showing that superficial PCs receive stronger SST+ interneuron inhibition when compared to deep PCs. This finding holds for each light pulse as there is no effect of light pulse on response ratio, one-way ANOVA: (F(2.418, 38.68) = 0.3875, p = 0.7199). **G)** Paired-pulse ratios are comparable between deep and superficial PCs; Paired t test (t(16) = 0.8643, p = 0.4002). N’s equivalent to (D). **H)** Summary diagram of circuit motifs between SST+ interneurons that target the proximal dendrites (e.g., bistratified cells) and pyramidal cells.

Next, for each paired recording between superficial and deep pyramidal cells we moved the focus of our 40X objective from the stratum radiatum to the stratum lacunosum moleculare (**Fig. 6A**). Strikingly, in same set of pyramidal cells, inhibitory synaptic inputs to the distal dendrites were of similar amplitudes (**Fig. 6B**). Indeed, across the population of paired recordings there was no bias in current amplitude (**Fig. 6C**). The evoked inhibitory currents were of small amplitude and depressed during the light pulse train, thus, we only analyzed responses to the first four light pulses. For the population, the average inhibitory current amplitude was the same in both superficial and deep pyramidal cells (**Fig. 6D**) and scaled consistently during the train (**Fig. 6E**). Finally, we found no difference in the paired pulse ratio (**Fig. 6F**), like that observed for input to the proximal dendrites (**Fig. 5G**). These data suggest OLM cells provide equal strength inhibition to the apical dendrites of both superficial and deep pyramidal cells (**Fig. 6G**).

**Figure 6.**
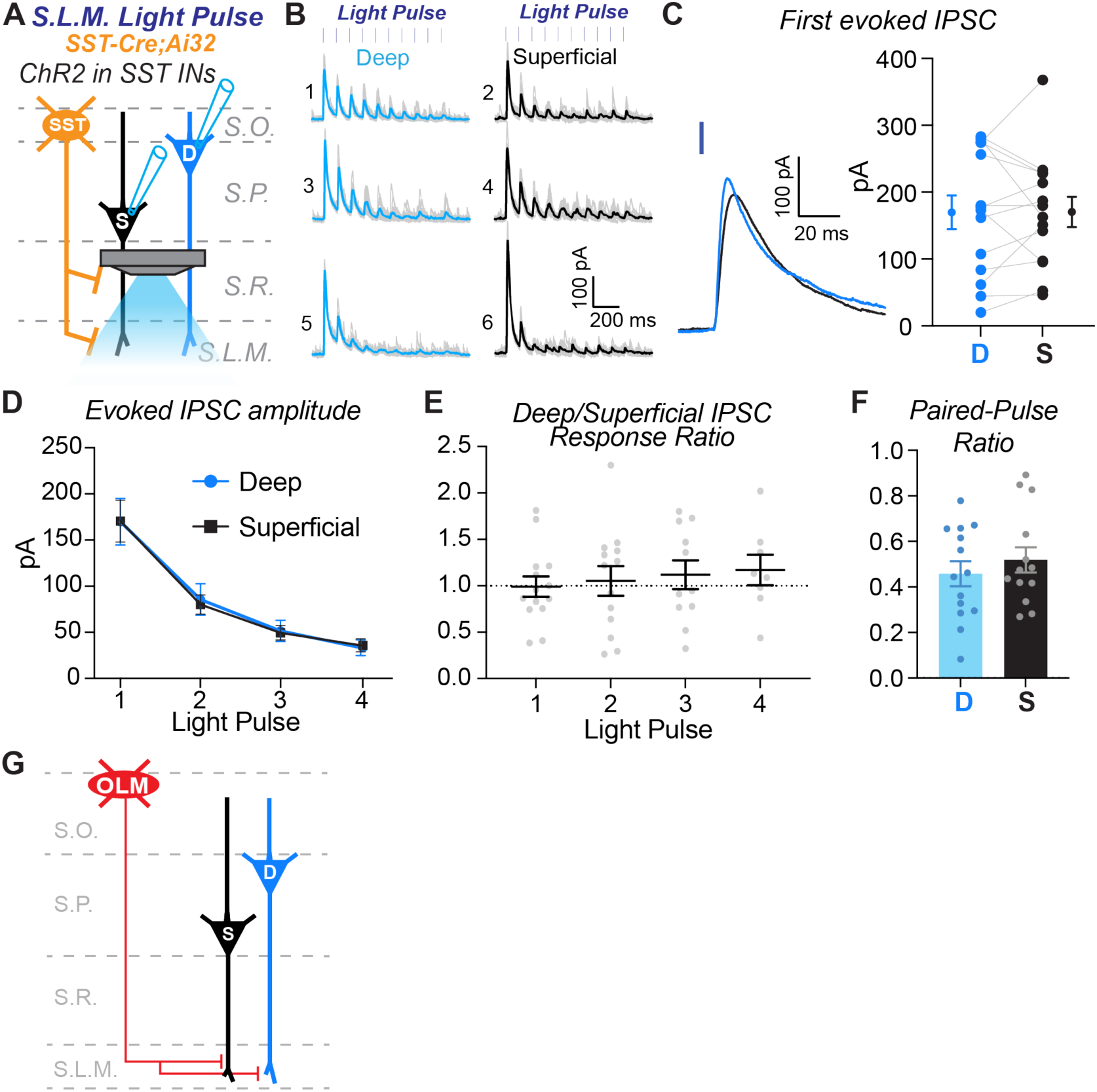
SST+ interneurons provide comparable inhibition to the distal dendrites of superficial pyramidal cells relative to deep pyramidal cells. **A)** Experimental configuration to assay the strength of inhibitory input from SST+ interneurons to the distal dendrites of superficial and deep pyramidal cells. Light was focused on the stratum lacunosum moleculare (S.L.M.) to excite SST+ interneuron axons. The same pairs from Figure 5 were recorded during light stimulation. **B)** ChR2-evoked inhibitory currents recorded from the same deep and superficial pyramidal cell pairs in Figure 5. Blue and black lines are the average responses from 10 trials; light grey lines are responses from individual trials. The numbers relate to the corresponding cell body locations in Figure 5B. **C)** Average amplitudes of responses to the first light pulse in the train for the population of recordings. Gray lines connect cells recorded as a pair. At left is an expanded example of ChR2-evoked responses to the first light pulse in a deep (blue) and superficial (black) pyramidal cell pair. n = 14 paired recordings from n = 3 mice. Paired t test (t(12) = 0.03988, p = 0.9688). **D)** Average ChR2-evoked current amplitudes for the population of responses recorded in deep and superficial PCs during the light pulse train. N’s equivalent to (C). No effect of cell type, repeated-measures two-way ANOVA: (F(1,13)= 0.0009285, p=0.9762). **E)** Ratios of the average peak current amplitudes between deep and superficial PCs for the population. Ratio of deep/superficial PC response to 10 optogenetic light pulses. Light pulse 1: n = 14 pairs; Light pulse 2, n = 13 pairs; Light pulse 3, n = 11 pairs; Light pulse 4, n = 9 pairs. The n’s are reduced for light pulses 2 – 4 because in a subset of paired recordings there was no evoked response in one of the cells. Dotted grey line is the assumed value if the ratio of the responses were equivalent. The average response ratios are at or near 1, showing that superficial and deep PCs receive comparable SST+ interneuron inhibition throughout the light pulse train. Mixed-effects one-way ANOVA: (F(2.362, 22.84) = 0.04451, p = 0.9723). **F)** Paired-pulse ratios are comparable between deep and superficial PCs. Deep PCs, n = 13; Superficial PCs, n = 14. Wilcoxon matched-pairs signed rank test, p = 0.7354. **G)** Summary diagram of circuit motif between SST+ OLM cells and pyramidal cells.

### OLMα2 cells provide greater inhibition to the distal dendrites of deep pyramidal cells compared to superficial pyramidal cells

By using SST-IRES-Cre;Ai32 mice to investigate inhibitory synapses to pyramidal cell apical dendrites we pooled input from OLM cells captured by the Nkx2.1-Cre and Chrna2-Cre lines. However, our data above indicate that Nkx2.1-Cre OLM cells and OLMα2 cells are separate subtypes that engage in unique circuits with superficial and deep pyramidal cells in CA1. Thus, we next generated Chrna2-Cre;Ai32 mice to directly investigate input from OLMα2 cells. As described above, we focused the excitation light on the stratum lacunosum moleculare and recorded inhibitory synaptic responses in pairs of superficial and deep pyramidal cells (**Figs. 7A and 7B**). In contrast to SST-IRES-Cre;Ai32 mice (**Fig. 6**), responses recorded in Chrna2-Cre;Ai32 mice were consistently larger in amplitude in deep compared to superficial pyramidal cells (**Fig. 7C**). Only the first response during the light pulse train was significantly larger (**Fig. 7D**), likely because the responses were small in amplitude and depressed during the train (**Figs. 7E and 7F**). However, the difference in amplitude was consistent across the paired recordings (**Fig. 7D**). Finally, there was no difference in paired pulse ratio (**Fig. 7G**). Combined with our data on excitatory connections from paired whole-cell recordings (**Figs. 2 – 4**), these data show that superficial pyramidal cells preferentially excite Nkx2.1-Cre+ OLM cells, which broadly inhibit the apical dendrites of all pyramidal cells (**Fig. 7H**). In contrast, OLMα2 cells preferentially inhibit the apical dendrites of deep pyramidal cells (**Fig. 7H**).

**Figure 7.**
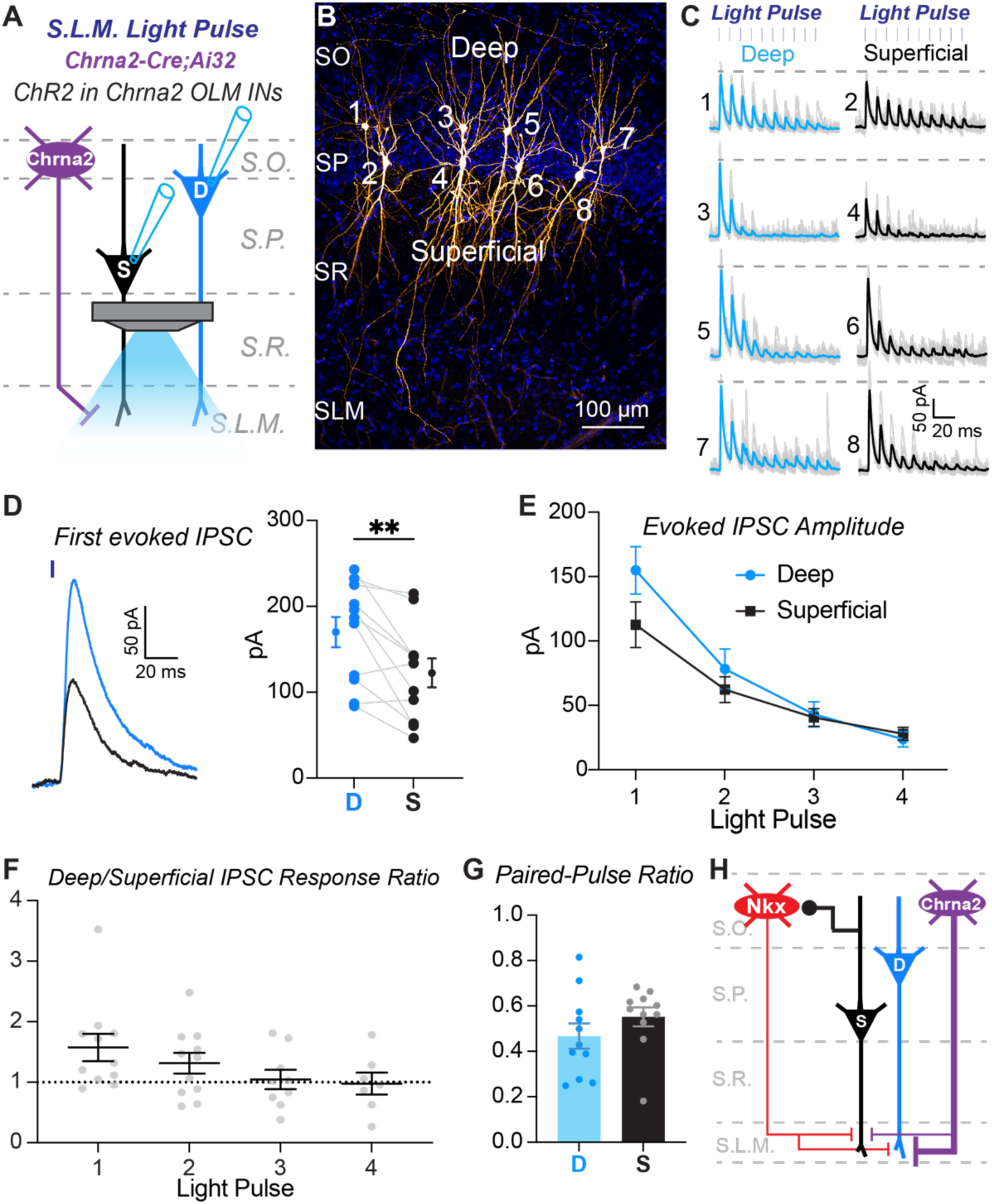
Chrna2+ OLM cells preferentially inhibit deep PCs compared to superficial PCs. **A)** Experimental configuration to assay the strength of inhibitory input from Chrna2-Cre+ interneurons to the distal dendrites of superficial and deep pyramidal cells. Light was focused on the stratum lacunosum moleculare (S.L.M.) to excite Chrna2-Cre+ interneuron axons. **B)** Representative example of a slice with recovered pyramidal cell bodies to demonstrate paired recordings of deep and superficial pyramidal cells in CA1. Four pairs were recorded in this slice (numbers 1+2, 3+4, 5+6, and 7+8). Note, the tissue was re-sectioned for cell recovery and slide mounting, thus the full dendritic tree is not shown in this section. **C)** ChR2-evoked inhibitory currents recorded from the deep and superficial pyramidal cell pairs shown in (B). Blue and black lines are the average responses from 10 trials; light grey lines are responses from individual trials. The numbers relate to the corresponding cell body locations in (B). Dashed gray line indicates the amplitude of the peak evoked response recorded in the deep PC in a pair. **D)** Average amplitudes of responses to the first light pulse in the train for the population of recordings. Gray lines connect cells recorded as a pair. At left is an expanded example of ChR2-evoked responses to the first light pulse in a deep (blue) and superficial (black) pyramidal cell pair. n = 11 paired recordings from n = 3 mice. ** p < 0.01, paired t test (t(10) = 3.174). **E)** Average ChR2-evoked current amplitudes for the population of responses recorded in deep and superficial PCs during the light pulse train. N’s equivalent to (D). Only the first response in the train is significantly different, thus there was no effect of cell type, repeated-measures two-way ANOVA: F(1,10) = 3.975, p = 0.0742. **F)** Ratios of the average peak current amplitudes between deep and superficial PCs for the population. Ratio of deep/superficial PC response to 10 optogenetic light pulses. Light pulse 1: n = 11 pairs; Light pulse 2, n = 11 pairs; Light pulse 3, n = 9 pairs; Light pulse 4, n = 7 pairs. The n’s are reduced for light pulses 2 – 4 because in a subset of paired recordings there was no evoked response in one of the cells. Dotted grey line is the assumed value if the ratio of the responses were equivalent. The average response ratios for light pulses 2 – 4 are at or near 1, showing that superficial and deep PCs receive comparable SST+ interneuron inhibition after the first light pulse in the train. Mixed-effects one-way ANOVA: (F(1.536, 12.29) = 2.460, p = 0.1341). **G**) Paired-pulse ratios are comparable between deep and superficial PCs; Wilcoxon matched-pairs signed rank test, p = 0.2402. N’s equivalent to (D). **H)** Summary diagram of circuit motif between Chrna2-Cre+ OLM cells and pyramidal cells versus Nkx2.1-Cre+ OLM cells and pyramidal cells.

### Biased inhibition of proximal dendrites depends on pyramidal cell radial position but not projection class

Pyramidal cells in CA1 are projection neurons that target diverse brain structures, and there is evidence that pyramidal cell projection class, in addition to radial position, can influence inhibitory circuit motifs. Specifically, in the deep layer, pyramidal cells that project to the amygdala receive stronger inhibition from PV+ basket cells compared to neighboring pyramidal cells that project to the medial prefrontal cortex (mPFC) (Lee et al., 2014). In the present study, the most striking inhibitory circuit motif we observed was that SST+ interneurons provide stronger inhibition to the proximal dendrites of superficial relative to deep pyramidal cells (**Fig. 5**). Thus, we next investigated if this circuit motif depends on pyramidal cell layer, projection class, or both. In SST-IRES-Cre;Ai32 mice, we injected green retrobeads into the mPFC and red retrobeads into the amygdala to differentially label pyramidal projection classes in CA1 (**Fig. 8A**). We first recorded pairs of mPFC- and amygdala-projecting pyramidal cells in the deep layer while delivering light pulses in the stratum radiatum (**Fig. 8B**). The ChR2-evoked inhibitory currents were equivalent between these projection classes (**Fig. 8C**), and there was no bias in current amplitude across the population of paired recordings. (**Fig. 8D**). Furthermore, throughout the light pulse train, the average amplitudes of inhibitory currents were not different (**Fig. 8E**). These data show that pyramidal cells belonging to different projection classes but in the same layer receive comparable inhibition along their proximal dendrites.

**Figure 8.**
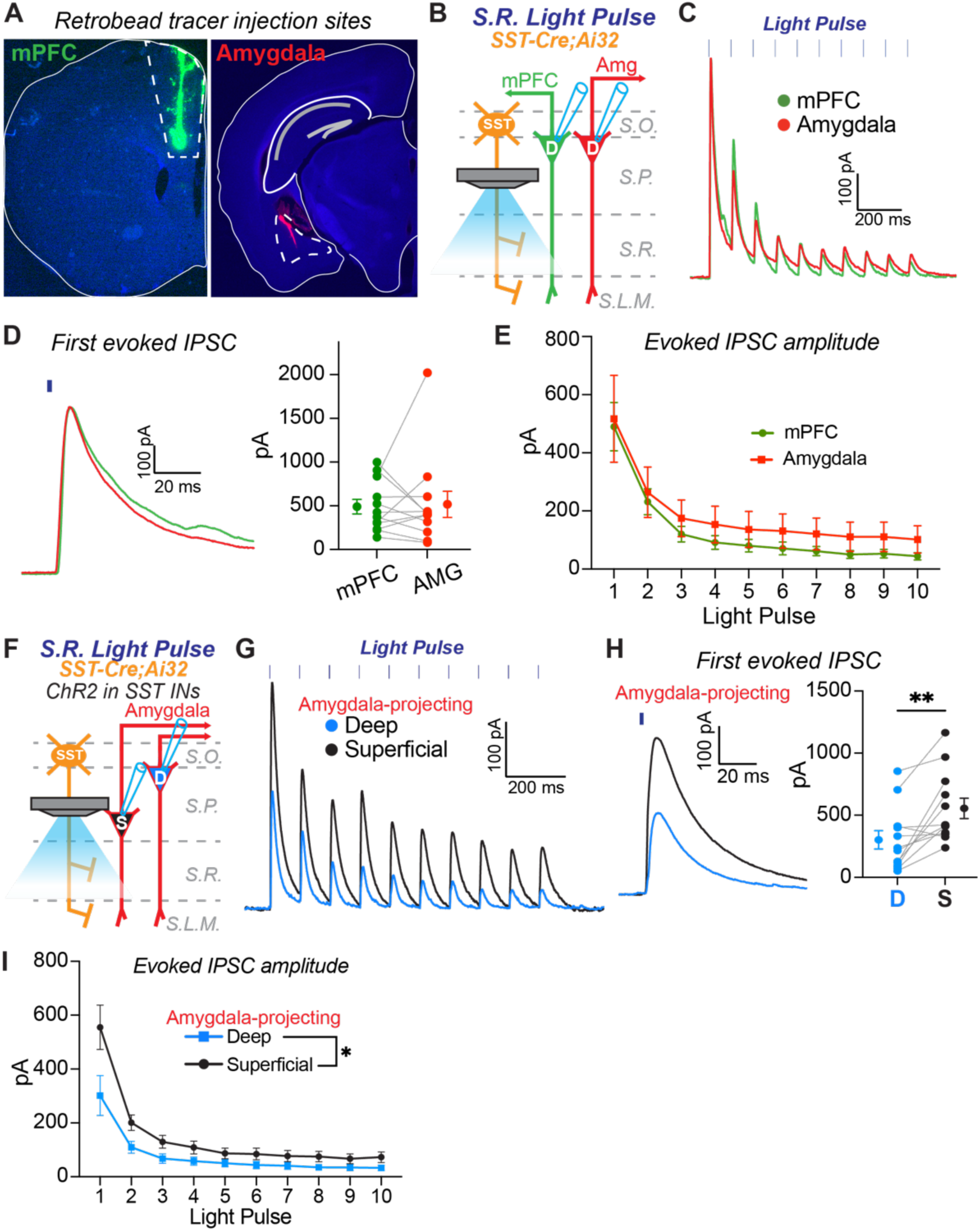
Pyramidal cell layer but not projection class determines biased inhibition to the proximal apical dendrites. **A)** Representative images of retrobead injections into medial prefrontal cortex (mPFC) and amygdala to label distinct pyramidal cell projection classes in CA1. Note, injections into the amygdala are at a 20-degree angle. **B)** Experimental configuration to assay the strength of inhibitory input from SST+ interneurons to the proximal dendrites of neighboring deep pyramidal cells belonging to two different projection classes. Light was focused on the stratum radiatum (S.R.) to excite SST+ interneuron axons. An mPFC-projecting and amygdala-projecting pyramidal cell were recorded simultaneously during light stimulation to directly compare inhibitory inputs. **C)** Representative example of average ChR2-evoked inhibitory currents recorded from neighboring mPFC-projecting (green) and amygdala-projecting (red) pyramidal cells in the deep layer. **D)** Average amplitudes of responses to the first light pulse in the train for the population of recordings. Gray lines connect cells recorded as a pair. At left is an expanded example of ChR2-evoked responses to the first light pulse in an mPFC-projecting deep pyramidal cell (green) and an amygdala-projecting deep pyramidal cell (red). n = 12 paired recordings from n = 3 mice. Wilcoxon matched-pairs signed rank test, p = 0.7910. **E)** Average ChR2-evoked current amplitudes for the population of responses during the light pulse train. N’s equivalent to (D). No effect of Projection Class, two-wav ANOVA: (F(1, 11)=1.188, p=0.2990). **F)** Experimental configuration to assay the strength of inhibitory input from SST+ interneurons to the proximal dendrites of deep and superficial pyramidal cells belonging to the same projection class. Light was focused on the stratum radiatum (S.R.) to excite SST+ interneuron axons. Deep and superficial amygdala-projecting pyramidal cells were recorded simultaneously during light stimulation to directly compare inhibitory inputs. **G)** Representative example of average ChR2-evoked inhibitory currents recorded from amygdala-projecting pyramidal cells in the deep (blue) and superficial (black) layers. **H)** Average amplitudes of responses to the first light pulse in the train for the population of recordings. Gray lines connect cells recorded as a pair. At left is an expanded example of ChR2-evoked responses to the first light pulse in a deep (blue) and superficial (black) pyramidal cell pair. n = 12 paired recordings from n = 3 mice. ** p < 0.01 Wilcoxon matched-pairs signed rank test. **I)** Average ChR2-evoked current amplitudes for the population of responses recorded in deep and superficial PCs during the light pulse train. N’s equivalent to (H). Main effect of layer, * p < 0.05, two-way ANOVA: (F(1, 11) = 8.668).

Next, we investigated if pyramidal cells belonging to the same projection class but in different layers receive differential dendritic inhibition. We recorded pairs of amygdala-projecting pyramidal cells in the superficial and deep layers while delivering light pulses in the stratum radiatum (**Fig. 8F**). Amygdala-projecting pyramidal cells in the superficial layer received stronger inhibitory inputs to their proximal dendrites than amygdala-projecting pyramidal cells in the deep layer (**Fig. 8G**). This bias was consistent across the population of paired recordings (**Fig. 8H**), resulting in larger inhibitory currents in superficial cells throughout the light pulse train (**Fig. 8I**).

Importantly, the results are consistent with our data from paired recordings of unlabeled pyramidal cell pairs in superficial and deep layers (**Fig. 5**). In summary, our data show that neighboring pyramidal cells in CA1 that project to the same target brain region, but are separated by layer, receive differential inhibitory input to their proximal dendrites. Collectively, our data suggest that pyramidal cells projecting to different targets, but in the same layer, are grouped by common inhibitory circuit motifs.

## DISCUSSION

SST+ interneurons include multiple cell-types that provide inhibition to distinct dendritic compartments of pyramidal cells to regulate their activity, synaptic integration, and synaptic plasticity (Müller and Remy, 2014; Pelkey et al., 2017; Fernández-Arroyo et al., 2024). Here, we show that in CA1 hippocampus SST+ interneurons form unique circuits with pyramidal cells based on the specific cell-type of both the pre- and postsynaptic neurons. These circuit motifs may contribute to the differential participation of deep and superficial pyramidal cells in network oscillations necessary for learning and memory and their differential encoding of spatial information during navigation (Klausberger and Somogyi, 2008; Valero et al., 2017; Soltesz and Losonczy, 2018; Topolnik and Tamboli, 2022).

We made the surprising finding that SST+ interneurons captured by the Nkx2.1-Cre and Htr3a-GFP lines, including both OLM cells and BiCs, engage in distinct circuits with deep and superficial pyramidal cells. These two transgenic lines are used to identify interneurons in the neocortex, hippocampus, and striatum that were derived from either the medial or caudal ganglionic eminence (MGE or CGE) during embryonic development (Xu et al., 2008; Lee et al., 2010). Mature interneurons are separated into major classes based on expression of molecular markers that include SST, PV, VIP, and CCK (Tremblay et al., 2016; Pelkey et al., 2017). These major interneuron classes are produced during embryogenesis primarily from the MGE and CGE; a small percent of interneurons is also generated in the preoptic area (POA) (Wonders and Anderson, 2006; Kepecs and Fishell, 2014; Kessaris et al., 2014; Pelkey et al., 2017). Progenitor cells in the MGE give rise to SST+ and PV+ interneuron classes, while those in the CGE give rise to the remainder, including VIP+ and CCK+ subtypes. Immature interneurons migrate out of the ganglionic eminences and disperse across the neocortex, hippocampus, and striatum. In the neocortex and hippocampus, there is very little overlap between cells captured by the Nkx2.1-Cre and Htr3a-GFP lines (Lee et al., 2010; Chittajallu et al., 2013). Concomitantly, SST+ interneurons in the neocortex are captured by the Nkx2.1-Cre line, but very rarely in Htr3a-GFP line (Lee et al., 2010). However, Chittajallu et al. (2013) found that approximately 30% of SST+ interneurons in the hippocampus, including both OLM cells and BiCs, are captured by the Htr3a-GFP line. They investigated Nkx2.1-Cre+ and Htr3a-GFP+ OLM cells in detail and found that their morphologies and electrophysiological properties were identical. However, during kainate-induced gamma oscillations in hippocampal slices, Htr3a-GFP+ OLM cells had lower firing rates and different phase preference than Nkx2.1-Cre+ OLM cells. Furthermore, Htr3a-GFP+ OLM cells responded to serotonergic agonists, but Nkx2.1-Cre+ OLM cells did not. Chittajallu et al. (2013) concluded that in the hippocampus there are two populations of SST+ cells that have separate embryonic origins and play different roles in regulating pyramidal cells. However, subsequent work challenged the conclusion that SST+ interneurons in the hippocampus originate from both the MGE and CGE (Harris et al., 2018; Asgarian et al., 2019; Winterer et al., 2019). Indeed, OLM cells captured by the Htr3a-Cre line (a companion BAC transgenic to the Htr3a-GFP line) express transcription factors associated with the MGE, including Lhx6, Satb1, and Sox6 (Winterer et al., 2019). Thus, OLM cells captured by the Nkx2.1-Cre and Htr3a-GFP lines likely both originate from the MGE. One possibility is that Htr3a-GFP+ OLM cells originate from progenitors in the dorsal MGE, which are captured by the Nkx6.2-Cre line, but not Nkx2.1-Cre line. Indeed Nkx2.1-Cre;Nkx6.2-Cre double transgenic mice capture almost all SST+ interneurons (Asgarian et al., 2019). However, overlap of expression in Htr3a-GFP;Nkx6.2-Cre mice throughout development has not been investigated.

Regardless of their embryonic origin, the existence of SST+ cells in the hippocampus that are captured selectively by the Htr3a-GFP or Nkx2.1-Cre mouse lines is puzzling and the extent to which they are functionally distinct has remained an open question. Our data extend and synthesize findings from previous work that show these two mouse lines indeed capture separate subsets of SST+ interneurons. We found that OLMα2 cells captured by the Chrna2-Cre line are a subset of Htr3a-GFP+ cells in the hippocampus. In a recent paper, Chamberland et al. (2024) showed that OLMα2 cells, relative to Nkx2.1-Cre+ OLM cells, preferentially inhibit PV+ interneurons. Thus, our data suggest that this circuit motif extends to Htr3a-GFP+ OLM cells. Furthermore, we identified new circuit motifs dependent on these mouse lines. First, SST+/Nkx2.1-Cre+ interneurons, but not OLMα2/Htr3a-GFP+ cells, receive biased excitatory synaptic input from superficial pyramidal cells compared to deep pyramidal cells. Second, OLMα2/Htr3a-GFP+ OLM cells preferentially inhibit deep pyramidal cells relative to superficial pyramidal cells. Collectively, these data show that the existence of SST+ interneurons that are nonoverlapping in Nkx2.1-Cre and Htr3a-GFP mice is not an artifact of these transgenic lines; these cells form distinct circuits and likely contribute to different functions in CA1.

We found that OLMα2 cells captured by the Chrna2-Cre line overlap with Htr3a-GFP+ cells in the hippocampus and subiculum but not in the neocortex. In neocortex, the Chrna2-Cre line captures a subset of layer 5 Martinotti cells (Hostetler et al., 2023; Wu et al., 2023; Chamberland et al., 2024), which are analogous to hippocampal OLM cells; they extend a long axon that ramifies in layer 1 at the distal tufts of pyramidal cell apical dendrites. Recent work found that there are multiple subtypes of Martinotti cells that express unique molecular markers, engage in unique circuits, and are captured by different transgenic lines (Munoz et al., 2017; Nigro et al., 2018; Hostetler et al., 2023; Wu et al., 2023; Agmon and Barth, 2024). Our work suggests a link between Htr3a-GFP+ hippocampal OLM cells and Chrna2-Cre+ neocortical Martinotti cells. It is possible that in the neocortex Htr3a-GFP expression is silenced during early development after Chrna2-Cre+ cells invade the cortex, due to cues from the local environment (Fishell and Kepecs, 2019). Indeed, in the hippocampus we observed a wide-range of GFP intensity in Chrna2-Cre-tdTomato+ cells; some were clearly GFP+ but dim (for examples, see **Figs. 1C and 1D**, comparing top and middle cells). Ultimately, it is unclear why neocortical Chrna2-Cre+ cells lack Htr3a-GFP expression. However, these data provide an interesting example for how similar interneuron cell-types can diverge between the hippocampus and neocortex.

Ultimately, our data can be combined with the existing literature to build a diagram of circuit motifs between inhibitory interneurons and deep and superficial pyramidal cells in CA1 (**Fig. 9A**). These circuit motifs suggest several mechanisms by which individual pyramidal cells or ensembles of cells might be coordinated in CA1 to support learning and memory or navigation (Mizuseki et al., 2011; Danielson et al., 2016; Geiller et al., 2017; Valero et al., 2017; Sharif et al., 2021). **1)** PV+ basket cells and OLMα2 cells may work in concert to preferentially regulate inhibition between the soma and apical dendrites of deep pyramidal cells (**Fig. 9B**). PV+ basket cells preferentially inhibit deep pyramidal cells to provide strong feedforward inhibition of excitatory input from CA3 (Lee et al., 2014; Masurkar et al., 2017; Hanson et al., 2025). Here, we found that OLMα2 cells preferentially inhibit the distal apical dendrites of deep pyramidal cells and thus may preferentially regulate excitatory input from the entorhinal cortex and thalamus. Furthermore, OLMα2 cells preferentially inhibit PV+ basket cells (Chamberland et al., 2024); thus, this may provide a mechanism for OLMα2 cells to selectively shift inhibition from the soma to the dendrites of deep pyramidal cells. **2)** We identified a novel feedback inhibitory circuit involving superficial pyramidal cells and dendrite-targeting interneurons (**Fig. 9C**). Superficial pyramidal cells provide biased excitatory drive to both Nkx2.1-Cre+ OLM and non-OLM cells (including BiCs) which could increase inhibition along the entire apical dendrite. Furthermore, interneurons that target the proximal dendrites in the stratum radiatum, which includes BiCs, provide biased inhibition to superficial pyramidal cells. This could serve to selectively regulate CA3 Schaffer collateral input to superficial cells. **3)** Our data suggest circuit mechanisms to allow pyramidal cells to communicate between layers, despite the lack of direct excitatory synaptic connections (**Fig. 9D**). For example, Chamberland et al. (2024) found that BiCs strongly inhibit PV+ basket cells. Thus, our data suggest that superficial pyramidal cells could preferentially excite BiCs to disinhibit deep pyramidal cells. Alternatively, superficial pyramidal cells could increase somatic inhibition of deep pyramidal cells via their biased direct excitation of PV+ basket cells (Lee et al., 2014; Hanson et al., 2025). Finally, superficial pyramidal cells could increase inhibition to the distal dendrites of deep pyramidal cells via strong excitatory drive to Nkx2.1-Cre+ OLM cells. New transgenic mouse lines and intersectional genetic approaches that target these specific interneuron subtypes will allow such circuits and their function to be directly investigated by future studies both in vitro and in vivo (Xu et al., 2008; Hostetler et al., 2023; Agmon and Barth, 2024; Chamberland et al., 2024).

**Figure 9.**
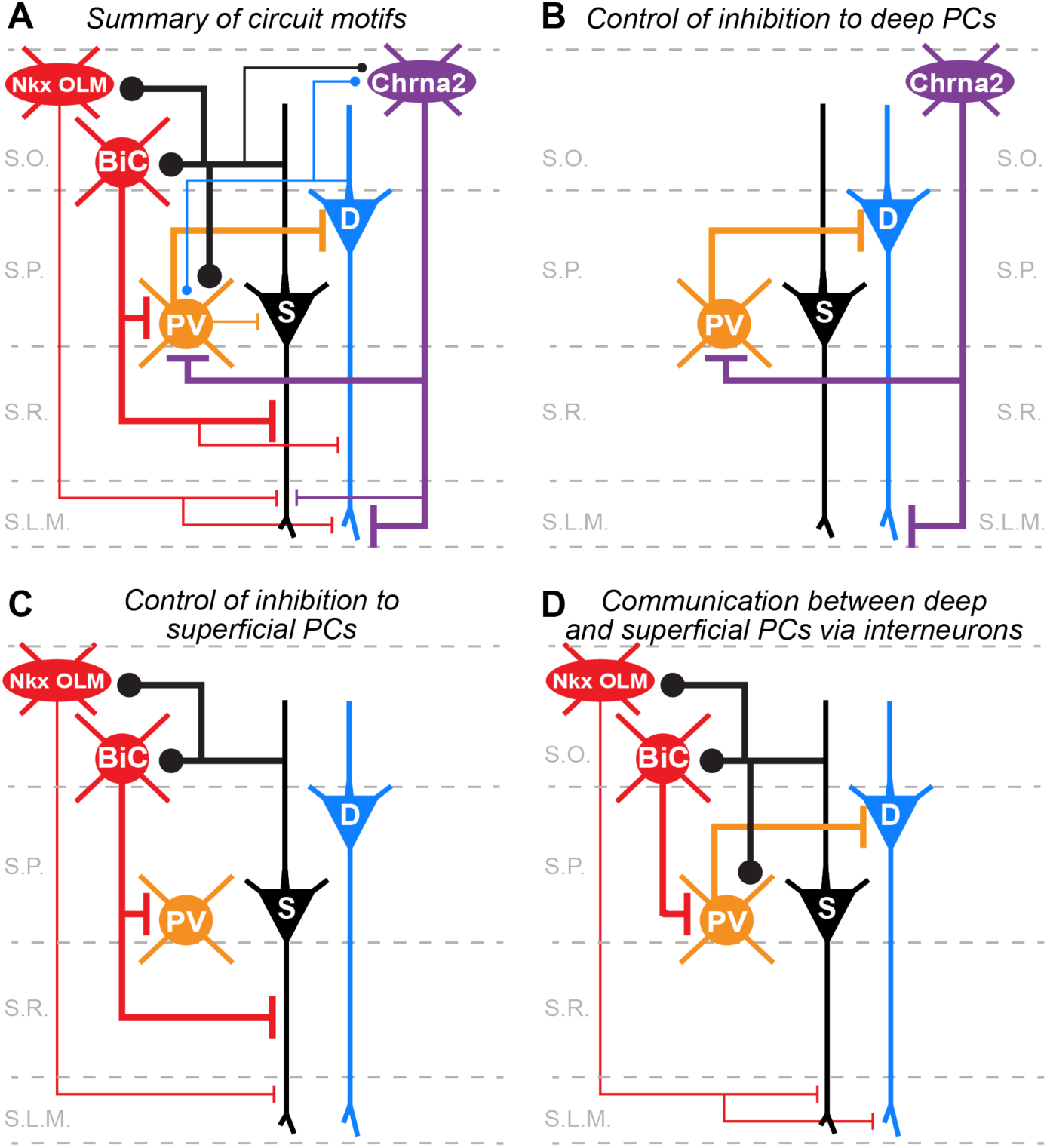
Summary of circuit motifs between pyramidal cells and interneurons in CA1. **A)** Summary of all synaptic connections and biases found in the present study and previous literature. Thicker lines indicate higher rates of synaptic connectivity observed in paired-whole cell recordings or larger amplitude synaptic responses observed in paired recordings or evoked using ChR2. **B)** PV+ basket cells and Chrna2-Cre+ OLM cells may work in concert to shift inhibition between the soma and apical dendrites of deep pyramidal cells. **C)** Superficial pyramidal cells engage in a feedback inhibitory circuit with both OLM and bistratified cells to that may shift inhibition between the soma and apical dendrites. **D)** Pyramidal cells in CA1 rarely form excitatory synaptic connections. However, biased connectivity with interneurons may provide a way for them to indirectly communicate via inhibition or disinhibition (see text for details).

## MATERIALS AND METHODS

### Animals

All experiments were conducted in accordance with animal protocols approved by the Ohio State University IACUC. Both male and female mice were used without bias. SST-IRES-Cre, Nkx2.1-Cre, Ai14, and Ai32 mice were obtained from the Jackson Laboratory (stock nos. 013044, 008661, 007914, and 012569). Chrna2-Cre mice (Leao et al., 2012) were obtained from the lab of Dr. Ariel Agmon (West Virginia University) with the permission of Dr. Klas Kullander (Uppsala University). Htr3a-GFP mice were obtained from Dr. Chris McBain (National Institute of Child Health and Human Development).

### Slice preparation

Mice (P21-P42 for paired whole-cell recordings or P33-P52 for optogenetics experiments) were anesthetized with isoflurane and then decapitated. The brain was dissected in ice-cold artificial cerebrospinal fluid (ACSF) containing (in mM): 100 sucrose, 80 NaCl, 3.5 KCl, 24 NaHCO_3_, 1.25 NaH_2_PO_4_, 4.5 MgCl, 0.5 CaCl_2_, and 10 glucose, saturated with 95% O_2_ and 5% CO_2_. Transverse sections of hippocampus were cut using a Leica VT 1200S vibratome (Leica Microsystems) and incubated in the above solution at 35 °C for 30 minutes post-dissection. For experiments to assay synaptic connections using paired recordings, we cut 300 μm coronal slices of dorsal CA1 hippocampus. For experiments to assay synaptic connections using optogenetics, we cut 400 μm horizontal sections of ventral CA1. Slices were then maintained at room temperature until use in recording ACSF containing (in mM): 130 NaCl, 3.5 KCl, 24 NaHCO_3_, 1.25 NaH_2_PO_4_, 1.5 MgCl, 2.5 CaCl_2_, and 10 glucose, saturated with 95% O_2_ and 5% CO_2_.

### Slice electrophysiology

For recording, slices were constantly perfused with ACSF at 2 mL/min at a temperature of 31-33 °C. Slices and cells were visualized using an upright microscope (Scientifica SliceScope) with a 40x water-immersion objective (Olympus), camera (Scientifica SciCam Pro), and Oculus software. Recordings were restricted to CA1. Deep and superficial pyramidal were identified by their proximity to the stratum oriens and stratum radiatum, respectively. Interneurons expressing tdTomato (via Cre recombination) or GFP, and pyramidal cells expressing green or red beads were identified by fluorescence (CoolLED pE-300ultra). Recording pipettes were pulled from borosilicate glass (World Precision Instruments) to a resistance of 3-5 MΩ using a vertical pipette puller (Narishige PC-100). Whole-cell patch clamp recordings were amplified using a Multiclamp 700B amplifier (Molecular Devices), filtered at 3 kHz (Bessel filter), and digitized at 20 kHz (Digidata 1550B and pClamp v11.1, Molecular Devices). Recordings were not corrected for liquid junction potential. Series resistance was closely monitored throughout recordings. Recordings were discarded if series resistance passed 25 MΩ.

For recordings of inhibitory interneurons to assay intrinsic membrane properties and excitatory postsynaptic currents in paired recording, the internal solution contained (in mM): 130 K-gluconate, 5 KCl, 2 NaCl, 4 MgATP, 0.3 NaGTP, 10 phosphocreatine, 10 HEPES, 0.5 EGTA, and 0.2% biocytin (Sigma-Aldrich catalog #B4261). The calculated ECl- for this solution was -79 mV. pH was adjusted to 7.4 with KOH. For recordings of pyramidal cells during paired recordings with inhibitory interneurons, the internal solution contained (in mM): 85 K-gluconate, 45 KCl, 2 NaCl, 4 MgATP, 0.3 NaGTP, 10 phosphocreatine, 10 HEPES, and 0.5 EGTA, for a calculated ECl- of -29 mV. pH was adjusted to 7.4 with KOH. We used this internal solution to allow switching between current and voltage clamp mode to test for unitary inhibitory synaptic connections. These connections were very rare and not analyzed. For experiments to record inhibitory currents in pyramidal cells using optogenetics, the internal solution contained (in mM): 135 CsMeSO_4_, 8 KCl, 4 MgATP, 0.3 NaGTP, 5 QX-314, 0.5 EGTA and 0.2% biocytin (Sigma-Aldrich catalog #B4261). pH was adjusted to 7.3 with CsOH.

For paired recordings of synaptic connections, presynaptic cells were recorded in current clamp and the membrane potential was biased to -70 mV. Presynaptic cells were made to fire trains of action potentials (25 at 50 Hz) using 2 nA steps of 2 millisecond duration every 10 seconds for 10 trials. Postsynaptic cells were recorded in voltage-clamp with a holding potential of -70 mV.

For optogenetic-assisted circuit mapping, we activated channelrhodopsin-2 (ChR2) by focusing blue light (CoolLED pE-300ultra, 460 nm) through a 40X objective centered over either the stratum radiatum or stratum lacunosum moleculare. Light pulses were of 2 millisecond duration and delivered in trains of 10 at 10 Hz every 10 seconds. During these recordings, 10 μM DNQX (Fisher, #23-121-0), and 50 μM DL-AP5 (Fisher, #36-931-0 R) were included in the bath to block AMPA and NMDA receptors. This was necessary to exclude potential excitatory drive from off-target Cre expression in pyramidal cells in SST-IRES-cre mice (**Fig. 1Ai**). Postsynaptic cells were recorded in voltage-clamp with a holding potential of 0 mV.

### Stereotaxic injections

For injections of retrograde tracer into the medial prefrontal cortex (mPFC) and amygdala, mice were anesthetized with 5% isoflurane and mounted in a stereotax (Neurostar, Germany). 24 hours prior to surgery, ibuprofen was added to the cage drinking water and maintained for 72 hours post-operation. Prior to surgery, mice were provided with extended release (48 – 72 hour) buprenorphine (1 mg/kg via subcutaneous injection) for additional post-operative analgesia. Red or green retrobeads (Lumafluor) were injected via a glass capillary nanoinjector (Neurostar, Germany) back filled with light mineral oil. The mPFC was targeted using the following coordinates: 1.86 mm rostral and -0.40 mm lateral to bregma, and 2.5 mm deep. The amygdala was targeted using the following coordinates: -1.90 mm caudal and -2.50 mm lateral to bregma, and 5.30 mm deep. For injections into the amygdala, the pipette was inserted at an angle of 20°. 100 nl of retrobeads were injected at 75 nl/min. The pipette was left in place for 10 min following the injection before removal. Mice were allowed to recover for at least two days before being sacrificed for experiments post-injection.

### Electrophysiology data analysis

All data were analyzed in Igor Pro v8.04 (WaveMetrics) using custom routines. pClamp files were imported into Igor using NeuroMatic (Rothman and Silver, 2018). Input resistance (Rin) was measured using a linear regression of voltage deflections (±20 mV from -70 mV resting membrane potential) in response to 1 s current steps. To calculate voltage sag, the membrane potential was biased to -70 mV (V_initial) followed by injection of a 1 second-duration negative current step of sufficient amplitude to reach a steady state Vm of -90 mV during the last 200 ms of the current injection (V_sag). The peak hyperpolarized Vm prior to sag (V_hyp) was used to calculate the sag index as (V_hyp – V_sag)/(V_hyp – V_initial). Capacitance was calculated as τ/Rin, both measured from the voltage response to a 10 pA current step relative to a -70 mV resting potential. The voltage threshold for action potential initiation was measured from the response to the minimum amplitude current step necessary to generate a spike. Spike threshold was defined as the voltage recorded at the maximum of the second derivative of voltage change with respect to time (Wilent and Contreras, 2005). Spike half-width at half-height was calculated between spike threshold and spike peak. Firing rate and adaptation ratio were collected from sweeps at twice threshold necessary to evoke an action potential. Adaptation ratio was calculated as the inter-spike interval between the first two action potentials divided by the averaged inter-spike interval between the last four action potentials.

To analyze postsynaptic currents in paired recordings, we first zeroed the data by subtracting the baseline and then performed 10 repetitions of binomial (Gaussian) smoothing. All postsynaptic currents (PSCs) were then analyzed relative to the timing of the peak of each presynaptic spike during the train. PSCs were detected by threshold crossing (-7 to -10 pA); if this threshold was already crossed at the instant of spike peak, the PSC data associated with that spike of that trial were discarded as being contaminated by spontaneous events. The proportion of failures was calculated as the ratio of evoked PSCs to the total number of non-contaminated trials for each presynaptic spike. The PSC potency was calculated as the average peak amplitude of all successfully evoked PSCs for each presynaptic spike (i.e., failures were not included in the average).

To analyze ChR2-evoked inhibitory postsynaptic currents (IPSCs) in pyramidal cell pairs, we first zeroed each sweep by subtracting the baseline (first 200 ms of each sweep prior to the first light pulse in a train). The IPSC average response was calculated as the average of the peak IPSC amplitude from 10 trials. To calculate the ratio of response amplitudes in deep versus superficial pyramidal cells in paired recordings, we used the average peak response to each light pulse. We required that average responses be of at least 10 pA, otherwise the ratio was not calculated.

### Principal Component Analysis and K-means clustering

These analyses were performed in RStudio. For principal component analysis (PCA) and K-means clustering analysis, we utilized up to 19 parameters from each interneuron including electrophysiology and morphological data (see **Figure 3**). These included Morphology, Bursting, Resting Vm, Rin, Sag, Tau, Capacitance, Spike1_time, Spike1_Threshold, Spike1_Height, Spike1_HalfWidth, Spike1_AHP, Spike2_time, Spike2_Threshold, Spike2_Height, Spike2_HalfWidth, Spike2_AHP, Firing Rate, Adaptation Ratio. Each parameter was normalized to the average value using the *.scale()* function. We then performed PCA using the *.prcomp()* function using the normalized data to determine which parameters best explain the variance between the interneurons. To identify clusters within the interneuron populations, we performed K-means analysis. First, we determined the optimal number of clusters via the Nbclust package in RStudio (min.nc was set to 2 and max.nc was set to 5). Next, we used the *kmeans()* function for K-means clustering using all normalized parameters. Finally, using the ggpubr package we generated a scatterplot based on the PCA and K-means clustering analysis to visualize the distribution of interneurons and their cluster assignments.

### Immunohistochemistry

To enhance the longevity of GFP expression in Chrna2-Cre;Ai14;Htr3a-GFP mice, we stained tissue sections from these mice with chicken anti-GFP (1:1000, Abcam catalog #ab13970, RRID:AB300798). Mice were anesthetized with ketamine/xylazine and tissue was fixed via transcardial perfusion with 4% paraformaldehyde. Brains were post-fixed for 1 hour in 4% paraformaldehyde. Brains were then cryopreserved in 30% sucrose, and then resectioned on a freezing microtome at 40 μm. Sections were rinsed in phosphate buffered saline (PBS), blocked for 2 hours in blocking buffer consisting of 10% normal goat serum with 0.5% Triton X-100 and 1% BSA, and then incubated in primary antibody (chicken anti-GFP) overnight at 4 °C. Sections were then rinsed with PBS and incubated in secondary antibody Alexa Fluor 488 Goat anti-chicken (1:1000, Invitrogen of Thermo Fisher Scientific catalog #A11039, Ref: A11039) and DAPI (1:2000, Thermo Fisher Scientific catalog #D1306) for 2 hours at room temperature or overnight at 4 °C. All antibodies were diluted in carrier solution containing PBS with 1% BSA, 1% normal goat serum, and 0.5% Triton X-100. Sections were then rinsed, mounted on slides, and cover slipped.

### Recovery of cell morphology

Following slice electrophysiology and biocytin cell filling (Sigma-Aldrich catalog #B4261), 300 μm tissue slices were drop-fixed for 24 hours in 4% paraformaldehyde. Slices were then rinsed with PBS and incubated in the secondary antibody streptavidin Alexa Fluor 488 (1:1000, Invitrogen of Thermo Fisher Scientific catalog #S311223), Alexa Fluor 555 (1:1000, Invitrogen of Thermo Fisher Scientific catalog #S32355), or Alexa Fluor 633 (1:1000, Invitrogen of Thermo Fisher Scientific catalog #S21375) and DAPI (1:2000, Thermo Fisher Scientific catalog #D1306) overnight at 4 °C. All antibodies were diluted in carrier solution with PBS with 1% BSA, 1% normal goat serum, and 0.5% Triton X-100. Slices were next rinsed with PBS and resectioned on a freezing microtome before being slide mounted. Slices containing interneurons were re-sectioned at 100 μm, while those containing pyramidal cells from optogenetic experiments were re-sectioned at 150 μm.

### Image acquisition and analysis

Tissue sections were imaged using a Leica DM6 CS confocal microscope with a 10X (for entire CA1 regions), 20X (for biocytin-filled cells and entire CA1) or 40X (for analysis of Chrna2+ and Htr3a co-expression in individual cells) objective. Images were processed in Fiji (Schindelin et al., 2012). Borders of each hippocampal strata (oriens, pyramidale, radiatum, and lacunosum moleculare) were determined by DAPI expression. Reconstructions were made using Simple Neurite Tracer (Longair et al., 2011) for Fiji (Schindelin et al., 2012). Tissue sections of retrobead injection sites were imaged using a QImaging Retiga R6 camera mounted to a Zeiss Axioskop2 microscope with a 1.25X objective.

### Statistical Analysis

Single-factor distributions were first tested for normality using a Shapiro-Wilk test. Distributions that did not violate normality were compared using a paired t test while those that violated normality were compared using the Wilcoxon matched pairs signed rank test. Data involving one factor were compared using a one-way ANOVA. Data involving two factors were compared using a repeated measures two-way ANOVA or a mixed-effects ANOVA. Connectivity rate was calculated using Fisher’s exact test. Each test and post hoc analysis used is described in the figure legend. Statistical significance was set at p < 0.05. Data in the text and graphs are reported as mean ± SEM. Statistical tests were performed using Graphpad Prism v10.1.1 (Graphpad Software, San Diego, CA).

## Author contributions

A.C.J. and J.C.W. designed the experiments. A.C.J., M.A.H., A.H.M., A.S., E.K.P., and N.B. collected the data. A.C.J. and J.C.W. analyzed the data. A.C.J. and J.C.W. wrote the paper.

## Acknowledgements

This work was supported by funding from the NIH (R01MH124870) to J.C.W. We thank Ariel Agmon (West Virginia University) for providing us with Chrna2-Cre mice with the permission of Klas Kullander (Uppsala University). We thank Chris McBain (NIH) for providing us with Htr3a-GFP mice.

